## Supplementary Data for "Going beyond size: Exploring the metabolic burden in *Pseudomonas putida* during heterologous protein production"

by

*Marleen Beentjes[1], Ana-Sofia Ortega-Arbulú[1], Carina Meiners, Andreas Kremling, Katharina Pflüger-Grau**

Professorship of Systems Biotechnology, Technical University of Munich, 85748 Garching (Germany)

[1] Both authors contributed equally to the work.

* For correspondence: Katharina Pflüger-Grau, Professorship of Systems Biotechnology, Technical University of Munich, Boltzmannstr. 15, 85748 Garching (Germany), Tel.: +49 89 289 15765; Fax.: +49 89 289 15766,

**Table T1: Oligonucleotides used in this work.** Restriction sites are shown in bold, the start codon is shaded in grey, and RBS is depicted in lowercase italics.

| Oligonucleotide | Sequence (5' – 3') | Description |
|---|---|---|
| noRBS-fw1 | GCGC**GAATTC**ATGAAAATCGAAGAAGGTAAAC | Forward primer for pSEVA-MBPeGFP_∆ |
| malE-fw_ceroni | ATAT**GAATTC***tactagagaaatcaaattaaggaggtaagata*ATGAAAATCGAAGAAGGT | Forward primer for pSEVA-MBPeGFP_str |
| SW20 | CCCCC**ACTAGT**TTAGTGATGATGATGATGATGGCTAGCCTTGTACAGCTCGTCCATGCC | Reverse primer for pSEVA-MBPeGFP_∆ and pSEVA-MBPeGFP_str and pSEVA-eGFP and pSEVA-eGFP_str |
| eGFP_fwd | CCCCC**GAATTC***aggaggaaaaacat*ATGGTGAGCAAGGGCGAGGA | Forward primer for pSEVA-eGFP |
| eGFP_str_fwd | CCCCC**GAATTC***tactagagaaatcaaattaaggaggtaagata*ATGGTGAGCAAGGGCGAGGA | Forward primer for pSEVA-eGFP_str |
| eGFP_rev | CCCCC**ACTAGT**TTACTTGTACAGCTCGTCCATGCC | Reverse primer for pSEVA-eGFP and pSEVA-eGFP_str |
| VioB_fwd | GCGC**GAATTC***aggaggaaaaacat*ATGAGCATTCTGGATTTCCCGC | Forward primer for amplification of *vioB* for assembly into pSEVA-VioBeGFP |
| VioB_rev | TGCTCACCATCGAACCACCACCACCGGCCTCGCGGCTCAGTTT | Reverse primer for amplification of *vioB* for assembly into pSEVA-VioBeGFP |
| eGFP_VioB_fwd | CCGCGAGGCCGGTGGTGGTGGTTCGATGGTGAGCAAGGGCGAG | Forward primer for amplification of *eGFP* for assembly into pSEVA-VioBeGFP |
| eGFP_VioB_rev | CTATCAACAGGAGTCCAAGATTAGTGATGATGATGATGATGGCTAG | Reverse primer for amplification of *eGFP* for assembly into pSEVA-VioBeGFP |
| fucI_fwd | GCCTAGGCCGCGGCCGCGCG**GAATTC***aggaggaaaaacat*ATGAAAAAAATCAGCTTACCG | Forward primer for amplification of *fucI* for assembly into pSEVA-FucIeGFP |
| fucI_rev | TGCTCACCATCGAACCACCACCACCACGCTTATACAACGGACC | Reverse primer for amplification of *fucI* for assembly into pSEVA-FucIeGFP |
| eGFP_fwd_fucI | GTATAAGCGTGGTGGTGGTGGTTCGATGGTGAGCAAGGGCGAG | Forward primer for amplification of eGFP for assembly into pSEVA-FucIeGFP |
| eGFP_rev_fucI | CTATCAACAGGAGTCCAAGA**ACTAG**TTTAGTGATGATGATGATGATGGCTAG | Reverse primer for amplification of eGFP for assembly into pSEVA-FucIeGFP |
| arsC (E. coli BL21)_fwd | GCCTAGGCCGCGGCCGCGCG**GAATTC***aggaggaaaaacat*ATGAGCAACATTACCATTTATCACAACC | Forward primer for amplification of *arsC* for assembly into pSEVA-ArsCeGFP |
| arsC (E. coli BL21)_rev | TGCTCACCATCGAACCACCACCACCTTTCAGGCGCTTACCCGC | Reverse primer for amplification of *arsC* for assembly into pSEVA-ArsCeGFP |
| eGFP _fwd_(arsC-BL21) | GCGCCTGAAAGGTGGTGGTGGTTCGATGGTGAGCAAGGGCGAG | Forward primer for amplification of eGFP for assembly into pSEVA-ArsCeGFP |

| | | |
|---|---|---|
| eGFP_rev_(arsC-BL21) | CTATCAACAGGAGTCCAAGA**ACTAG**TTTAGTGATGATGATGATGATGGCTAG | Reverse primer for amplification of eGFP for assembly into pSEVA-ArsCeGFP |
| arsCeGFP_StrongRBS_fwd | ATAT**GAATTC***tactagagaaatcaaattaaggaggtaagata*ATGAGCAACATTACCATTTATCACAAC | Forward primer for pSEVA-ArsCeGFP_str |
| Standard_fwd | CAGGCGAAGAGATCCTG | Forward primer for qPCR |
| Standard_rev | GATGAAGCGACGTGCCTC | Reverse primer for qPCR |
| Fusion_proteins_fwd | GTGGTGGTGGTTCGATG | Forward primer for qPCR |
| Fusion_proteins_rev | TCAGCTTGCCGTAGGTG | Reverse primer for qPCR |

**Table T2: Plasmids used in this work.**

| Plasmid | Description | Reference |
|---|---|---|
| pSEVA438 | Empty plasmid | (*1*) |
| pSEVA-MBPeGFP | Standard RBS, MBPeGFP production | (*2*) |
| pSEVA-MBPeGFP_Δ | No RBS, MBPeGFP production | This work |
| pSEVA-MBPeGFP_str | Strong RBS, MBPeGFP production | This work |
| pSEVA-eGFP | Standard RBS, eGFP production | This work |
| pSEVA-eGFP_str | Strong RBS, eGFP production | This work |
| pSEVA-ArsCeGFP | Standard RBS, ArsCeGFP production | This work |
| pSEVA-FucIeGFP | Standard RBS, FucIeGFP production | This work |
| pSEVA-VioBeGFP | Standard RBS, VioBeGFP production | This work |
| pSEVA-ArsCeGFP_str | Strong RBS, ArsCeGFP production | This work |

**Table T3: Composition of the amino acid mixture used for supplementation of the medium.** Note that no pure L-tyrosine was included (due to solubility issues), and that L-leucine, L-phenylalanine, and L-tryptophan have lower concentrations**.**

| Amino acid | Concentration [mM] |
|---|---|
| L-alanine | 5 |
| L-arginine | 5 |
| L-aspargine | 5 |
| L-aspartic acid | 5 |
| L-cysteine | 5 |
| L-glutamic acid | 5 |
| L-glutamine | 5 |
| L-glycine | 5 |
| L-glycyl-l-tyrosine | 5 |
| L-histidine | 5 |
| L-isoleucine | 5 |
| L-lysine | 5 |
| L-methionine | 5 |
| L-proline | 5 |
| L-serine | 5 |
| L-threonine | 5 |
| L-valine | 5 |
| L-leucine | 2.5 |
| L-phenylalanine | 1 |
| L-tryptophan | 1 |

**Table T4: Predicted translation initiation rates.**

| Protein | Predicted translation rate * | | Fold change |
|---|---|---|---|
| | Standard RBS | Strong RBS | |
| MBPeGFP | 27048.90 | 515482.77 | 19 |
| ArsCeGFP | 10848.81 | 22831.93 | 2 |

*The translation initiation rate was calculated from the mRNA sequence directly downstream of the *Pm* promoter to the *SpeI* restriction site using the RBS calculator (predict mode Version v2.1.1) (*3*)

A

| Percentage [%] | MBPeGFP | ArsCeGFP | FucIeGFP | VioBeGFP |
|---|---|---|---|---|
| **Ala** | 8,50 | 4,00 | 8,70 | 10,10 |
| **Arg** | 1,90 | 4,30 | 4,50 | 7,80 |
| **Asn** | 5,50 | 4,80 | 4,40 | 3,00 |
| **Asp** | 6,70 | 6,50 | 5,90 | 6,90 |
| **Cys** | 0,30 | 1,00 | 1,70 | 1,00 |
| **Gln** | 2,70 | 2,50 | 3,40 | 3,60 |
| **Glu** | 6,90 | 7,50 | 6,60 | 5,90 |
| **Gly** | 8,80 | 8,80 | 9,40 | 8,80 |
| **His** | 2,90 | 4,50 | 3,70 | 3,70 |
| **Ile** | 5,60 | 6,00 | 5,60 | 3,00 |
| **Leu** | 8,20 | 9,30 | 7,40 | 11,40 |
| **Lys** | 9,10 | 7,00 | 5,30 | 2,60 |
| **Met** | 2,10 | 2,80 | 3,30 | 2,70 |
| **Phe** | 4,30 | 3,80 | 3,40 | 4,50 |
| **Pro** | 5,00 | 5,00 | 3,90 | 6,50 |
| **Ser** | 3,90 | 4,50 | 5,60 | 5,00 |
| **Thr** | 5,80 | 6,80 | 5,80 | 3,80 |
| **Trp** | 1,40 | 0,30 | 1,80 | 1,50 |
| **Tyr** | 4,20 | 3,50 | 3,30 | 3,50 |
| **Val** | 6,10 | 7,00 | 6,30 | 4,80 |

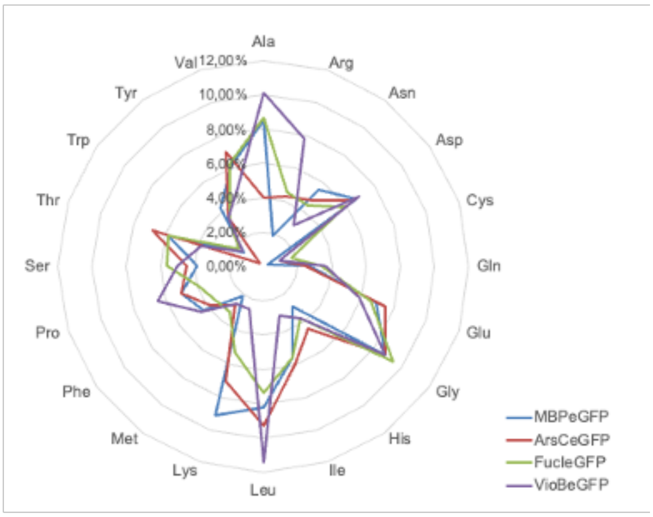


B

| Count | MBPeGFP | ArsCeGFP | FucIeGFP | VioBeGFP |
|---|---|---|---|---|
| **Ala** | 53 | 16 | 73 | 126 |
| **Arg** | 12 | 17 | 38 | 97 |
| **Asn** | 34 | 19 | 37 | 37 |
| **Asp** | 42 | 26 | 50 | 86 |
| **Cys** | 2 | 4 | 14 | 13 |
| **Gln** | 17 | 10 | 29 | 45 |
| **Glu** | 43 | 30 | 56 | 74 |
| **Gly** | 55 | 35 | 79 | 110 |
| **His** | 18 | 18 | 31 | 46 |
| **Ile** | 35 | 24 | 47 | 38 |
| **Leu** | 51 | 37 | 62 | 142 |
| **Lys** | 57 | 28 | 45 | 33 |
| **Met** | 13 | 11 | 28 | 34 |
| **Phe** | 27 | 15 | 29 | 56 |
| **Pro** | 31 | 20 | 33 | 81 |
| **Ser** | 24 | 18 | 47 | 62 |
| **Thr** | 36 | 27 | 49 | 47 |
| **Trp** | 9 | 1 | 15 | 19 |
| **Tyr** | 26 | 14 | 28 | 44 |
| **Val** | 38 | 28 | 53 | 60 |
| | 69 kD | 47 kD | 93 kD | 139 kD |

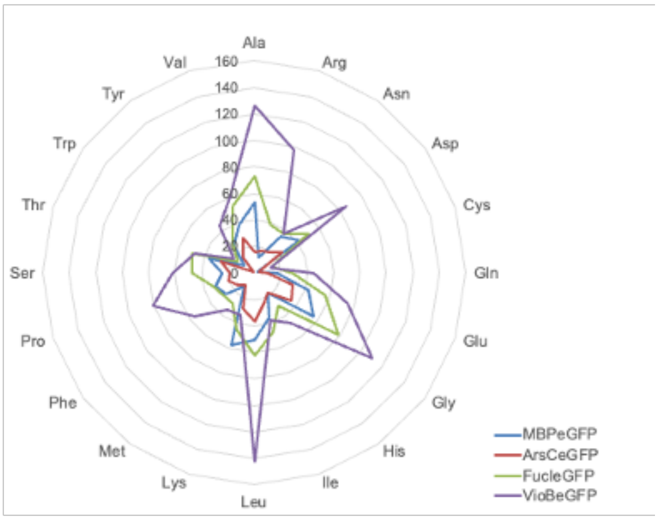


**Figure S1: Overview of the size and amino acid composition of the four alternative eGFP fusion proteins. A** Percentage of amino acids per protein molecule and **B** absolute number of amino acids per protein molecule and size of the different fusion proteins in kDa.

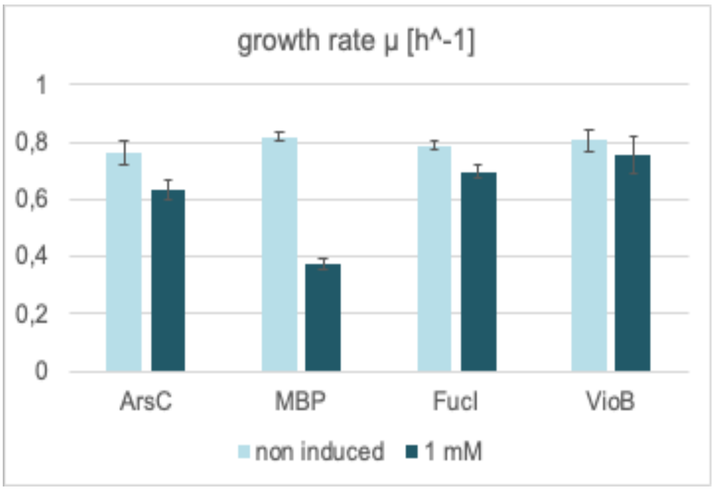


**Figure S2: Growth rate of *P. putida CAP* producing the alternative fusion proteins ArsCeGFP, MBPeGFP, FucIeGFP, or VioBeGFP**. *P. putida* CAP carrying the respective plasmid was grown in M9 medium with glucose (3 g/L) in a 96-well microtiter plate, and gene expression was induced with 1 mM 3-MB; experiments were performed in biological triplicate.

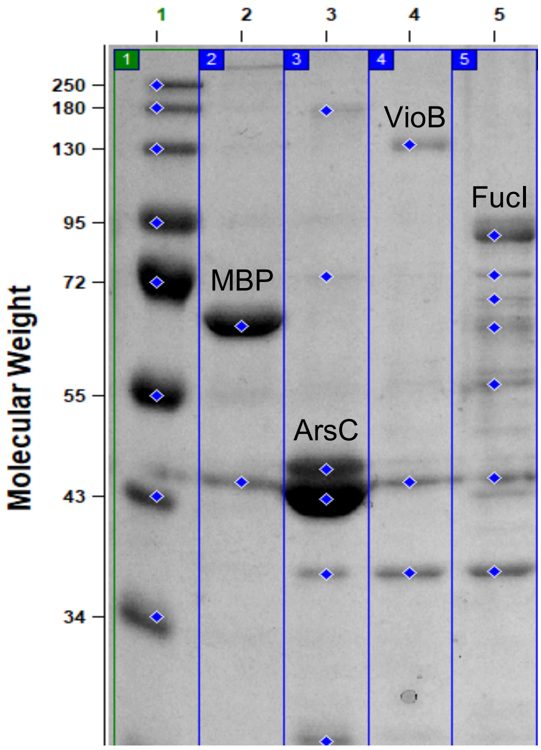


**Figure S3: SDS-PAGE of purified fusion proteins**. Proteins were purified via the C-terminal His tag from a 50 mL culture at OD 2.0 in M9 minimal medium (10 g $L^{-1}$ glucose), induction with 1 mM 3-MB, and separated on a 12% SDS-PAGE according to their size. Note that all Fusion protein match their expected size (MBPeGFP 69 kDa; ArsCeGFP 47 kDa; VioBeGFP 139 kDa; FucIeGFP 93 kDa).

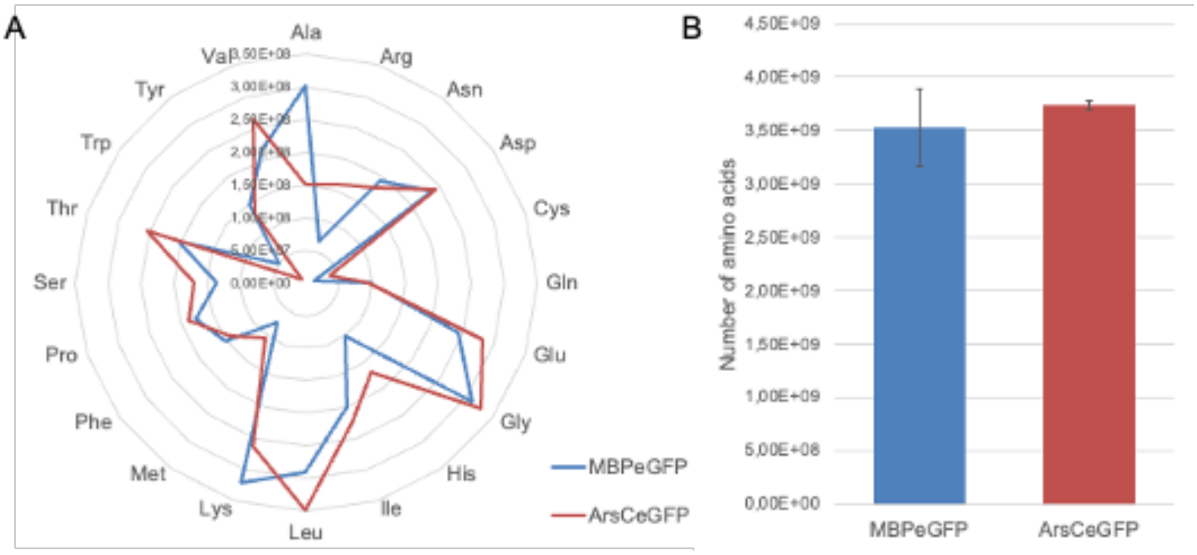


**Figure S4: Distribution (A) and number of amino acids (B) needed for the production of the amount of heterologous protein per cell.** The number of produced protein molecules of each of the indicated fusion proteins per cell at an OD=2 of the respective culture was estimated assuming 5.04*$10^8$ cells per ml at OD=1 (*4*).

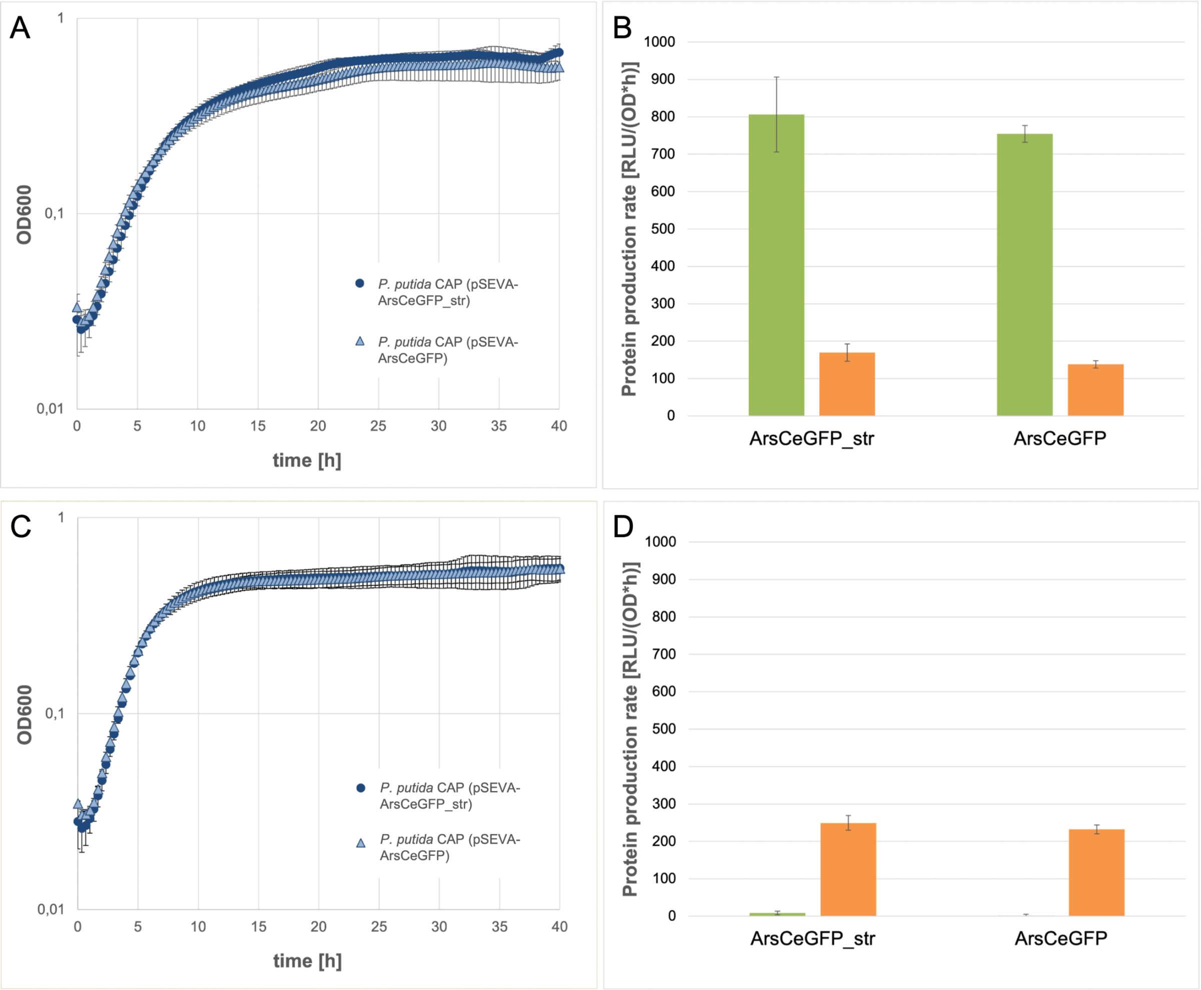


**Figure S5: Effect of a stronger ribosome binding site on growth and protein production rates of the alternative burden protein ArsC.** Growth of *P. putida* CAP (pSEVA-ArsCeGFP_str) (dark blue circles) and *P. putida* CAP (pSEVA-ArsCeGFP) (blue triangles) and protein production rates of the burden protein ArsCeGFP (green) and the capacity monitor mCherry (orange) in inducing (**A&B**) or non-inducing (**C&D**) conditions. <u>*Cultivation conditions*</u>: *M9 medium with glucose (3 g/L), 96-well microtiter plate, cultivation at 30 °C: induction with 1 mM 3-MB (A&B) or without 3-MB addition (C&D); experiments were performed in biological triplicates.*

### Bibliography

1. E. Martínez-García, S. Fraile, E. Algar, T. Aparicio, E. Velázquez, B. Calles, H. Tas, B. Blázquez, B. Martín, C. Prieto, L. Sánchez-Sampedro, M. H. H. Nørholm, D. C. Volke, N. T. Wirth, P. Dvořák, L. Alejaldre, L. Grozinger, M. Crowther, A. Goñi-Moreno, P. I. Nikel, J. Nogales, V. de Lorenzo, SEVA 4.0: an update of the Standard European Vector Architecture database for advanced analysis and programming of bacterial phenotypes. *Nucleic Acids Res.* **51**, D1558–D1567 (2022).

2. P. Vogeleer, P. Millard, A.-S. O. Arbulú, K. Pflüger-Grau, A. Kremling, F. Létisse, Metabolic impact of heterologous protein production in *Pseudomonas putida*: Insights into carbon and energy flux control. *Metab. Eng.* **81**, 26–37 (2024).

3. H. M. Salis, The ribosome binding site calculator. *Methods in Enzymology* **498**, 19–42 (2011).

4. P. Mira, P. Yeh, B. G. Hall, Estimating microbial population data from optical density. *PLoS ONE* **17**, e0276040 (2022).
